# Towards creating an extended metabolic model (EMM) for *E. coli* using enzyme promiscuity prediction and metabolomics data

**DOI:** 10.1101/536060

**Authors:** Sara A. Amin, Elizabeth Chavez, Vladimir Porokhin, Nikhil U. Nair, Soha Hassoun

**Affiliations:** Department of Computer Science, Tufts University, Medford, MA; Department of Biology, University of North Carolina, Chapel Hill, NC; Department of Chemical and Biological Engineering, Tufts University, Medford, MA; Department of Computer Science and Department of Chemical & Biological Engineering, Tufts University, Medford, MA

**Keywords:** Metabolic engineering, enzyme promiscuity, extended metabolic model, systems biology, enzyme activity prediction

## Abstract

**Background:** Metabolic models are indispensable in guiding cellular engineering and in advancing our understanding of systems biology. As not all enzymatic activities are fully known and/or annotated, metabolic models remain incomplete, resulting in suboptimal computational analysis and leading to unexpected experimental results. We posit that one major source of unaccounted metabolism is promiscuous enzymatic activity. It is now well-accepted that most, if not all, enzymes are promiscuous – i.e., they transform substrates other than their primary substrate. However, there have been no systematic analyses of genome-scale metabolic models to predict putative reactions and/or metabolites that arise from enzyme promiscuity.

**Results:** Our workflow utilizes PROXIMAL – a tool that uses reactant-product transformation patterns from the KEGG database – to predict putative structural modifications due to promiscuous enzymes. Using iML1515 as a model system, we first utilized a computational workflow, referred to as Extended Metabolite Model Annotation (EMMA), to predict promiscuous reactions catalyzed, and metabolites produced, by natively encoded enzymes in *E. coli*. We predict hundreds of new metabolites that can be used to augment iML1515. We then validated our method by comparing predicted metabolites with the *Escherichia coli* Metabolome Database (ECMDB).

**Conclusions:** We utilized EMMA to augment the iML1515 metabolic model to more fully reflect cellular metabolic activity. This workflow uses enzyme promiscuity as basis to predict hundreds of reactions and metabolites that may exist in *E. coli* but may have not been documented in iML1515 or other databases. We provide detailed analysis of 23 predicted reactions and 16 associated metabolites. Interestingly, nine of these metabolites, which are in ECMDB, have not previously been documented in any other *E. coli* databases. Four of the predicted reactions provide putative transformations parallel to those already in iML1515. We suggest adding predicted metabolites and reactions to iML1515 to create an Extended Metabolic Model (EMM) for *E. coli*.

## Background

The engineering of metabolic networks has enabled the production of high-volume commodity chemicals such as biopolymers and fuels, therapeutics, and specialty products [1–5]. Producing such compounds requires transforming microorganisms into efficient cellular factories [6–9]. Biological engineering has been aided via computational tools for constructing synthesis pathways, strain optimization, elementary flux mode analysis, discovery of hierarchical networked modules that elucidate function and cellular organization, and many others (e.g., [10–14]). These design tools rely on organism-specific metabolic models that represent cellular reactions and their substrates and products. Model reconstruction tools [15, 16] use homology search to assign function to Open Reading Frames obtained through sequencing and annotation. Once the function is identified, the corresponding biochemical transformation is assigned to the gene. Additional biological information such as gene-protein-reaction associations is utilized to refine the models. Exponential growth in sequencing has resulted in an “astronomical”, or better yet, “genomical”, number of sequenced organisms [17]. There are now databases (e.g., KEGG [18], BioCyc [19], and BiGG [20]) that catalogue organism-specific metabolic models. Despite progress in sequencing and model reconstruction, the complete characterizing of cellular activity remains elusive, and metabolic models remain incomplete. One major source of uncatalogued cellular activity is attributed to orphan genes. Because of limitations of homology-based prediction of protein function, there are millions of protein sequences that are not assigned reliable functions [21]. Integrated strategies that utilize structural biology, computational biology, and molecular enzymology continue to address assigning function to orphan genes [22].

We focus in this paper on another major source of uncatalogued cellular activity − promiscuous enzymatic activity, which has recently been referred to as ‘underground metabolism’ [23–25]. While enzymes have widely been held as highly-specific catalysts that only transform their annotated substrate to product, recent studies show that enzymatic promiscuity – enzymes catalyzing reactions other than their main reactions – is not an exception but can be a secondary task for enzymes [26–31]. More than two-fifths (44%) of KEGG enzymes are associated with more than one reaction [32]. Promiscuous activities however are not easily detectable *in vivo* since, i) metabolites produced due to enzyme promiscuity may be unknown, ii) product concentration due to promiscuous activity may be low, iii) there is no high-throughput way to relate formed products to specific enzymes, and iv) it is difficult to identify potentially unknown metabolites in complex biological samples. Outside of *in vitro* biochemical characterization studies to predict promiscuous activities, there are few resources that record details about promiscuous enzymes such as MINEs Database [33], and ATLAS [34]. Despite the current wide-spread acceptance of enzyme promiscuity, and its prominent utilization to engineer catalyzing enzymes in metabolic engineering practice [35–38], promiscuous enzymatic activity is not currently fully documented in metabolic models. Advances in computing and the ability to collect large sets of metabolomics data through untargeted metabolomics provide an exciting opportunity to develop methods to identify promiscuous reactions, their catalyzing enzymes, and their products that are specific to the sample under study. The identified reactions can then be used to complete existing metabolic models.

We describe in this paper a computational workflow that aims to extend preexisting models with reactions catalyzed by promiscuous native enzymes and validate the outcomes using published metabolomics datasets. We refer to the augmented models as Extended Metabolic Models (EMMs), and to the workflow to create them as EMMA (EMM Annotation). Each metabolic model is assumed to have a set of reactions and their compounds and KEGG reaction IDs. Each reaction, and thus transformation, is assumed to be reversible unless indicated otherwise. EMMA utilizes PROXIMAL [39], a method for creating biotransformation operators from KEGG reactions IDs using RDM (Reaction Center, Difference Region, and Matched Region) patterns [40], and then applying the operators to given molecules. While initially developed to investigate products of Cytochrome P450 (CYP) enzymes, highly promiscuous enzymes utilized for detoxification, the PROXIMAL method is generic. To create an EMM for a known metabolic model, PROXIMAL generates biotransformation operators for each reaction in the model and then applies the operators to known metabolites within the model. The outcome of our workflow is a list of putative metabolites due to promiscuous enzymatic activity and their catalyzing enzymes and reactions. In this work, we apply EMMA to iML1515, a genome-scale model of *Escherichia coli* MG1655 [41]. EMMA predicts hundreds of putative reactions and their products due to promiscuous activities in *E. coli*. The putative products are then compared to measured metabolites as reported in *Escherichia coli* Metabolome Database, ECMDB [42, 43]. We identify 23 new reactions and 16 new metabolites that we recommend adding to the *E. coli* model iML1515. Four of these reactions have not been catalogued prior for *E. coli* or other organisms, suggesting novel undocumented promiscuous transformations, while five other reactions are catalogued for species other than *E. coli.* Further, there were ten reactions that were cataloged in other *E. coli* databases (e.g. EcoCyc [44], and KEGG), but not in iML1515. These 19 reactions led to the addition of the 16 metabolites that are new to iML1515. Additionally, there were four new reactions that present putative transformation routes that are in parallel to existing reactions in *E. coli.* No new metabolites are added due to these four reactions.

## Results

The application of PROXIMAL to iML1515 yielded a lookup table with 1,875 biotransformation operator entries. The operators were applied on two sets of metabolites. One set consisted of 106 iML1515 metabolites with predicted or measured concentration values above 1 μM [45]. We focused on these metabolites as the assumption is that high concentration metabolites are more likely to undergo transformation by promiscuous enzymatic activity and form detectible derivatives. When applied to this set, the operators predicted the formation of 1,423 known (with PubChem IDs) metabolites of which 57 were identified to exist in *E. coli* per ECMDB. After manual curation (per Step 1 in the Methods section), our workflow recommended 16 new metabolites and 23 reactions that can be used to augment the iML1515 model. The second set of metabolites consisted of the non-high concentration metabolites in iML1515. Our workflow predicted the formation of 3,694 known (with PubChem IDs) metabolites. Out of the predicted metabolites of the second set 210 derivatives are found in ECMDB. We provide a listing of all derivatives in **Supplementary File 1**. For the remainder of the Results section, we focus on detailed analysis of derivative products due to high-concentration metabolites. Results of Flux Balance Analysis and Flux Variability Analysis for the added EMMA reactions are reported in **Supplementary File 2**.

Identified reactions were divided into four categories, C1–C4. The rationale for the various categories is explained using a decision tree (**Fig. 1**), a machine learning model that classifies data into groupings that share similar attributes [46]. With the exception of leaf nodes, each node in the tree tests the presence or absence of a particular attribute. Left branches represent the presence of the attribute, while the right branch represents the attribute’s absence. Each leaf node represents a classification category and is associated with a subset of the 23 reactions. At the root node of the decision tree, we tested if a PROXIMAL predicted metabolite is in the iML1515 model. If it is, and if the enzyme catalyzing the reaction within iML1515 model producing this metabolite is different than the enzyme PROXIMAL used to predict the relevant biotransformation, then it is classified in Category 1 (C1). Reactions belonging to C1 are parallel transformation to the ones in the model. They represent novel biotransformation routes between existing metabolites since they are generated using a different gene/enzyme than what is reported in iML1515. If previous conditions do not apply to the predicted product, then it is discarded as the reaction is already in iML1515.

**Fig. 1:**
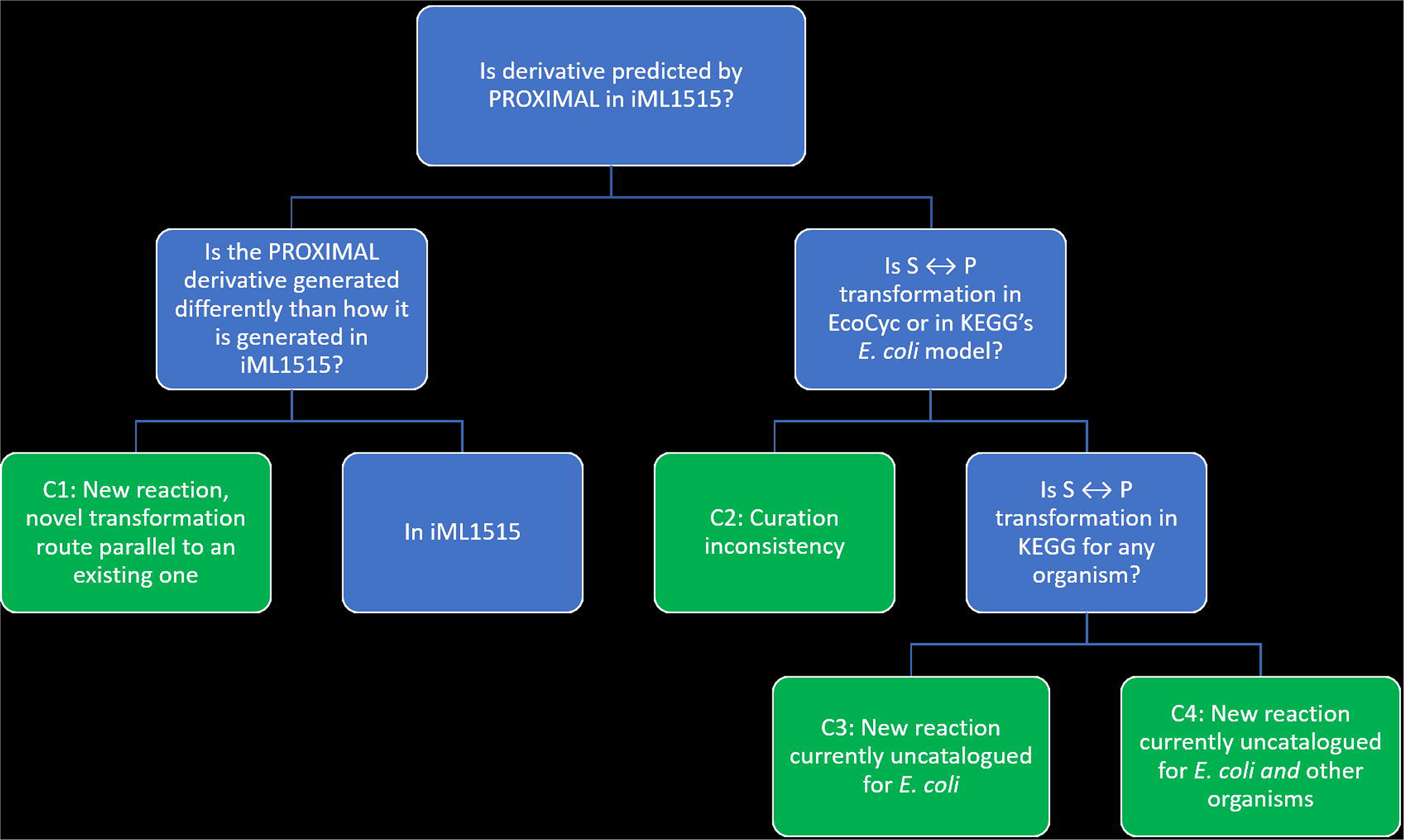
Decision tree for classifying reactions identified based on enzyme promiscuity. When analyzing the iML1515 *E. coli* model, reaction categories C1, C2, C3, and C4 had 4, 10, 5, and 4 predicted reactions, respectively.

If a predicted metabolite is not one of the known metabolites in iML1515, the decision tree determines whether the predicted metabolite and reaction set is associated with *E. coli* in other databases (KEGG and EcoCyc). If the biotransformation is present in KEGG or EcoCyc, then the predicted metabolite is classified into Category 2 (C2), reflecting a curation issue where some reactions were not included in the iML1515 model. If the predicted metabolite is not in iML1515 and not associated with *E. coli* in KEGG nor listed in EcoCyc, then the decision tree determines if the same chemical transformation (same substrate and same product) is documented to occur in other organisms. Predicted biotransformations documented in KEGG for organisms other than *E. coli* are classified in Category 3 (C3). While biotransformations not found in KEGG are classified as Category 4 (C4).

Each Category consists of a set of reactions. C1 consists of four reactions that are predicted to be catalyzed by enzymes that are different than those in iML1515. The details of the predicted reactions are shown in **Fig 2**, and **Table 1** details a comparison between those predicted reactions and their parallel reactions in iML1515. The phosphoribosyltransferase reaction between cytosine and cytidine-5’-monophosphate (CMP) is predicted to occur in *E. coli* due to EC 2.4.2.10 (orotate phosphoribosyltransferase) (**Fig. 2A**) and that between 2-oxoglutarate and 2-hydroxyglutarate by EC 1.1.1.79 (glyoxylate reductase) (**Fig. 2B**). We also predict the transformation between bicarbonate and carboxyphosphate catalyzed by EC 3.6.1.7 (acylphosphatase) (**Fig. 2C**). While carboxyphosphate is not in iML1515, the transformation is considered parallel to a reaction catalyzed by EC 6.3.5.5 that is documented to occur for *E. coli* in KEGG (see **Fig. 3J**). The last prediction is the coenzyme A transferase reaction between acetoacetyl-CoA and acetoacetate due to EC 2.8.3.10 (citrate CoA-transferase) (**Fig. 2D**).

**Table 1:**
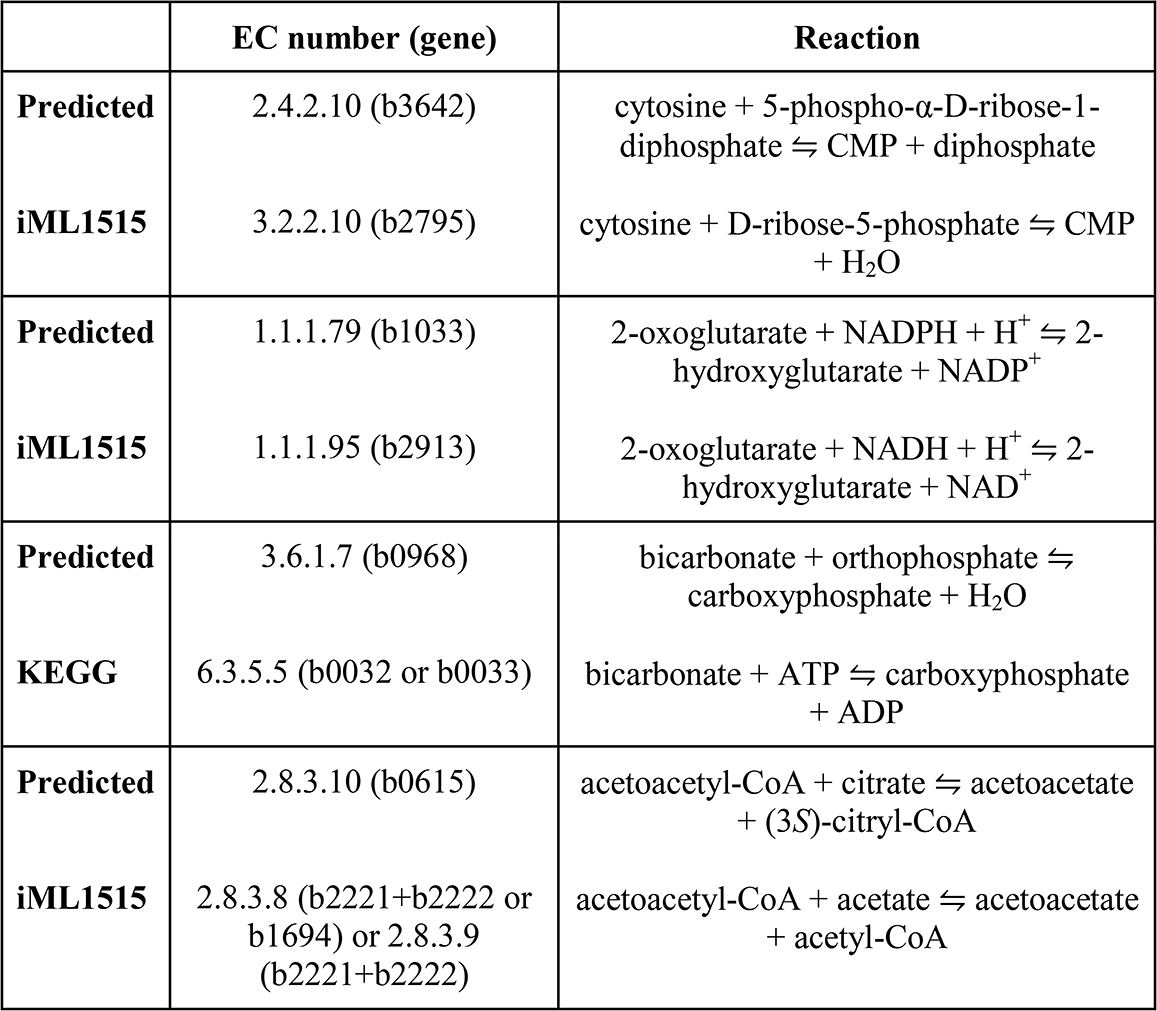
List of C1 reactions predicted by EMMA and their parallel reactions in *E. coli* iML1515. Each of the Predicted/iML1515 reaction pair occurs between the same substrate and product but utilize different co-substrate or cofactors.

|  | EC number (gene) | Reaction |
| --- | --- | --- |
| **Predicted** | 2.4.2.10 (b3642) | cytosine + 5-phospho-α-D-ribose-1-diphosphate ⇌ CMP + diphosphate |
| **iML1515** | 3.2.2.10 (b2795) | cytosine + D-ribose-5-phosphate ⇌ CMP + H<sub>2</sub>O |
| **Predicted** | 1.1.1.79 (b1033) | 2-oxoglutarate + NADPH + H<sup>+</sup> ⇌ 2-hydroxyglutarate + NADP<sup>+</sup> |
| **iML1515** | 1.1.1.95 (b2913) | 2-oxoglutarate + NADH + H<sup>+</sup> ⇌ 2-hydroxyglutarate + NAD<sup>+</sup> |
| **Predicted** | 3.6.1.7 (b0968) | bicarbonate + orthophosphate ⇌ carboxyphosphate + H<sub>2</sub>O |
| **KEGG** | 6.3.5.5 (b0032 or b0033) | bicarbonate + ATP ⇌ carboxyphosphate + ADP |
| **Predicted** | 2.8.3.10 (b0615) | acetoacetyl-CoA + citrate ⇌ acetoacetate + (3S)-citryl-CoA |
| **iML1515** | 2.8.3.8 (b2221+b2222 or b1694) or 2.8.3.9 (b2221+b2222) | acetoacetyl-CoA + acetate ⇌ acetoacetate + acetyl-CoA |

**Fig. 2:**
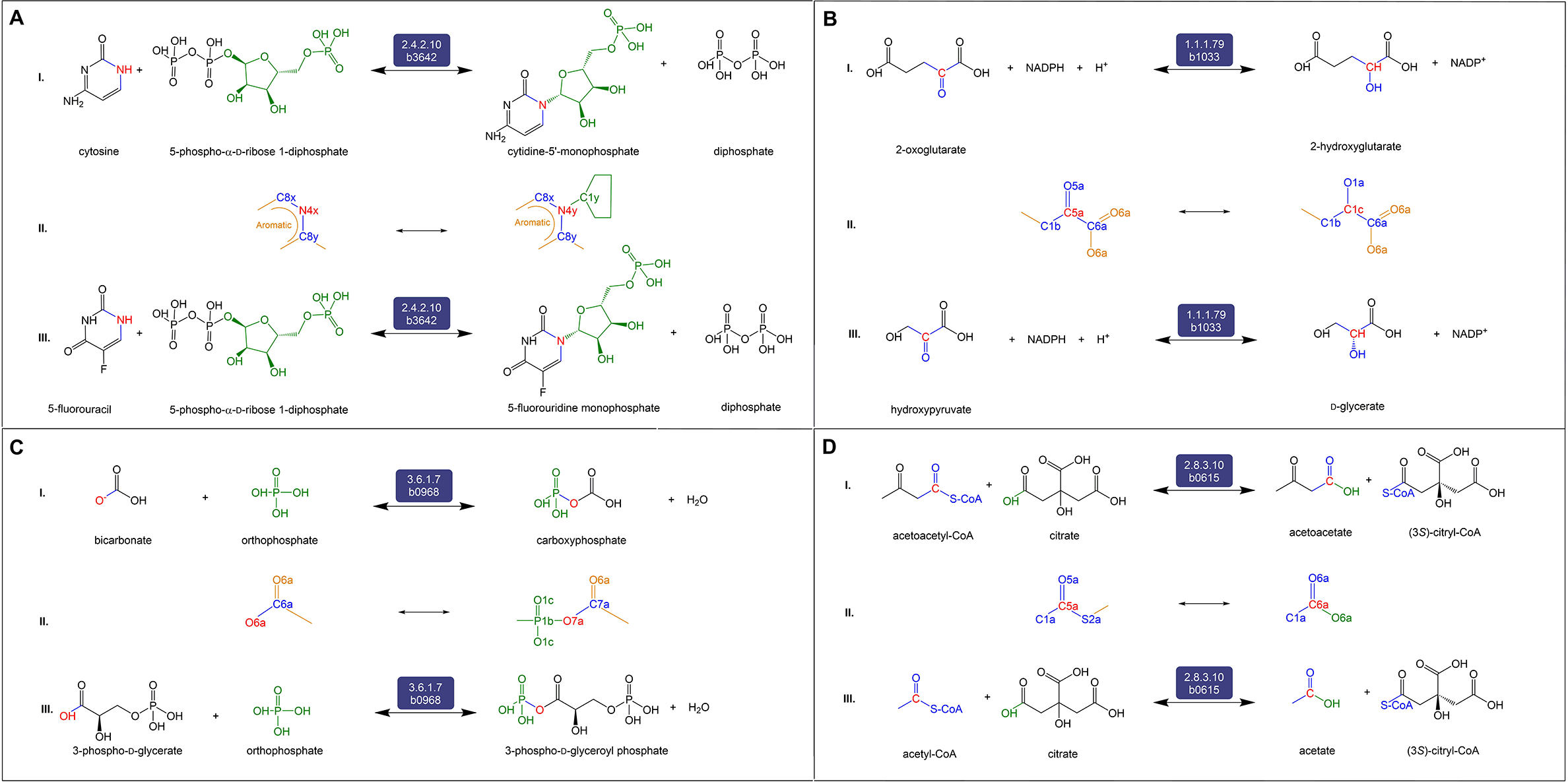
The set of four reactions belonging to Category 1 (C1). Reactions in C1 are predicted to be catalyzed by enzymes different than those in iML1515. Each of the four panels is divided into three sections I) the balanced reaction developed by our workflow indicating the reactants, products, and the promiscuous enzyme, II) the RDM pattern showing the Reaction Center (R) in red where the biotransformation occurs, and III) the native reaction catalyzed by the potentially promiscuous enzyme, as catalogued in KEGG.

**Fig. 3:**
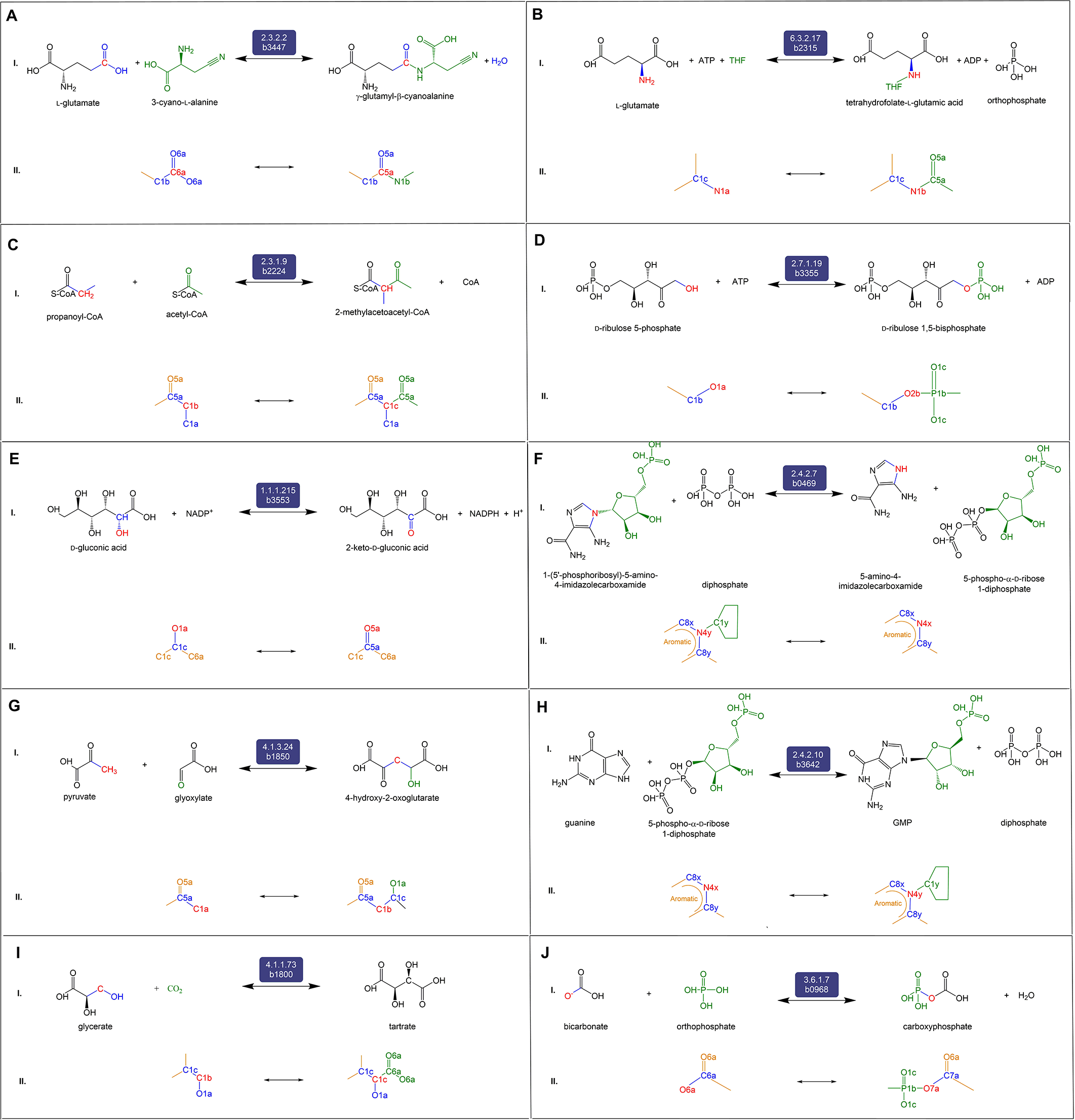
The set of ten reactions belonging to Category 2 (C2). Reactions in C2 are associated with derivatives not present in iML1515 but are associated with *E. coli* in KEGG and/or EcoCyc. Each of the ten panels is divided into two sections I) the balanced reaction developed by our workflow, that is also documented in KEGG, indicating the reactants, products, and the promiscuous enzyme, and II) the RDM pattern showing the Reaction Center (R) in red where the biotransformation occurs.

C2 consists of 10 reactions known to be in *E. coli* but missing from the iML1515 model. The first predicted reaction is the aminoacyltransferase reaction between L-glutamate and *γ*-glutamyl-β-cyanoalanine due to EC 2.3.2.2 (*γ*-glutamyltransferase) (**Fig. 3A**). The second is a predicted ligase reaction between L-glutamic acid and THF to form/consume THF-L-glutamic acid by EC 6.3.2.17 (tetrahydrofolate synthase) (**Fig. 3B**). The third is an acyltransferase transformation between propanoyl-CoA and 2-methylacetoacetyl-CoA catalyzed by EC 2.3.1.9 (acetoacetyl-CoA thiolase) (**Fig. 3C**). Fourth, PROXIMAL predicted the phosphotransferase reaction between of D-ribulose-5-phosphate and D-ribulose-1,5-bisphosphate by EC 2.7.1.19 (phosphoribulokinase) (**Fig. 3D**). The fifth predicted reaction known to be in *E. coli* is the redox transformation of D-gluconic acid to 2-keto-D-gluconic acid by EC 1.1.1.215 (gluconate 2-dehydrogenase) (**Fig. 3E**). The workflow also predicted glycosyltransferase transformation of 5-amino-4-imidazolecarboxamide to/from 1-(5’-phosphoribosyl)-5-amino-4-imidazolecarboxamide by EC 2.4.2.7 (AMP pyrophosphorylase) (**Fig. 3F**). The seventh predicted reaction is the transformation between pyruvate and 4-hydroxy-2-oxoglutarate by EC 4.1.3.24 (**Fig. 3G**). The eighth reaction is catalyzed by EC 2.4.2.10 to transform guanine to/from GMP (**Fig. 3H**). Also, PROXIMAL predicted the transformation between glycerate and tartrate by EC 4.1.1.73 (**Fig. 3I**). Lastly, bicarbonate is transformed to/from carboxyphosphate by EC 3.6.1.7 (**Fig. 3J**).

C3 consists of five predicted reactions that are not documented in *E. coli* but are known in other organisms. The first of these, the transformation between pyruvate and 4-carboxy-4-hydroxy-2-oxoadipate (**Fig. 4A**) catalyzed by EC 4.1.3.17 (HMG aldolase), is present in many organisms, including bacteria, as part of the benzoate degradation pathway (KEGG R00350). The transformation is predicted to occur in *E. coli* due to EC 4.1.3.34 (citryl-CoA lyase). Both EC 4.1.3.17 and EC 4.1.3.34 are lyases enzymes that form carbon-carbon bonds. 4-Carboxy-4-hydroxy-2-oxoadipate is known to be formed/consumed by EC 4.2.1.80 (2-keto-4-pentenoate hydratase) in *E. coli* (KEGG R04781). Another predicted reaction is the (de)aminating redox transformation between L-histidine and imidazol-5-yl-pyruvate, catalyzed by EC 1.4.1.4 (glutamate dehydrogenase) (**Fig. 4B**). Imidazol-5-yl-pyruvate is not known to be produced in any other way in *E. coli*, according to ECMDB and KEGG databases. The transformation of L-histidine to/from imidazol-5-yl-pyruvate is known to occur in the bacterium *Delftia acidovorans* by EC 2.6.1.38 (histidine transaminase) [47]. C3 also includes the predicted aryltransferase reaction between geranyl diphosphate and geranyl hydroxybenzoate by EC 2.5.1.39 (4-hydroxybenzoate transferase) (**Fig. 4C**). While the general reaction of all-*trans*-polyprenyl diphosphate to 4-hydroxy-3-polyprenylbenzoate is known to occur in *E. coli*, the specific transformation between geranyl diphosphate to geranyl hydroxybenzoate is known to occur in plants as part of shikonin biosynthesis, by EC 2.5.1.93 (4-hydroxybenzoate geranyltransferase) [48]. The fourth predicted reaction is the redox transformation between D-alanine and 2-aminoacrylic acid (**Fig. 4D**). This reaction is predicted to be catalyzed by EC 1.3.1.98 (UDP-*N*-acetylmuramate dehydrogenase). While 2-aminoacrylic acid is not known to be produced in *E. coli* in any other way, the transformation between D-alanine and 2-aminoacrylic acid occurs in other organisms such as *Staphylococcus aureus* [49]. Lastly, our workflow predicts the transformation between phenylpyruvate and phenyllactate by EC 1.1.1.100 (**Fig. 4E**). This transformation is known to occur in plants by EC 1.1.1.237 [50].

**Fig. 4:**
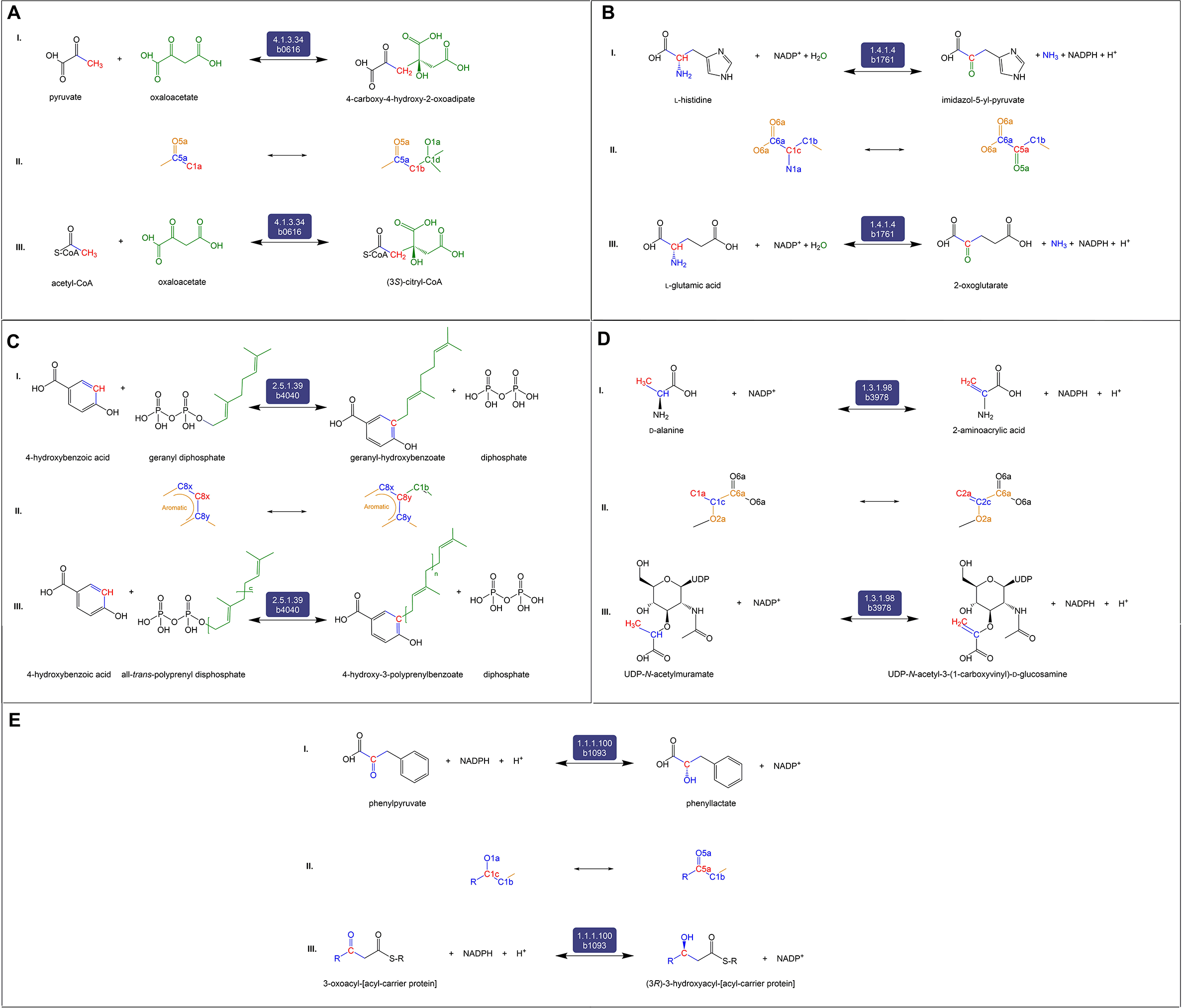
The set of five reactions belonging to Category 3 (C3). C3 reactions and derivatives are neither present in iML1515 nor associated with *E. coli* in KEGG and EcoCyc. However, according to KEGG, the reactions occur in other organisms. Each of the five panels is divided into three sections I) the balanced reaction developed by our workflow indicating the reactants, products, and the promiscuous enzyme, II) the RDM pattern showing the Reaction Center (R) in red, and III) the native reaction catalyzed by the potentially promiscuous enzyme, as catalogued in KEGG.

C4 consists of four predicted reactions that are not currently catalogued in KEGG for any organism (**Fig. 5**). The first reaction (**Fig. 5A**) is the oxidoreductive interconversion between aminomalonate and L-serine by EC 1.1.1.23 (histidinol dehydrogenase). There is one reaction (KEGG R02970) catalyzed by EC 2.6.1.47 (L-alanine:oxomalonate aminotransferase) that produces aminomalonate; but it is not a redox reaction and is associated with rat and silkworm, not *E. coli* [51]. The second is a hydrolytic decarboxylation reaction between *N*-acetylputrescine and *N*-acetylornithine (**Fig. 5B**) predicted to be catalyzed by EC 4.1.1.36 (PPC decarboxylase). The product, *N*-acetylputrescine, is involved in a number of enzymatic reactions – ECs 1.4.3.4 (monoamine oxidase), 2.3.1.57 (spermidine acetyltransferase), and 3.5.1.62 (acetylputrescine deacetylase) – in many organisms that include both eukaryotes and bacteria [16]. The third reaction in this category is the hydrolytic decarboxylation reaction between 3-ureidopropionate and *N*-carbamoyl-L-aspartate also catalyzed by EC 4.1.1.36 (PPC decarboxylase). 3-Ureidopropionate is present in eukaryotes and bacteria (but not *E. coli*) and is involved in reactions catalyzed by ECs 3.5.1.6 (β-ureidopropionase) and 3.5.2.2 (dihydropyrimidinase). The last reaction is the transformation between D-gluconic acid and D-galactarate by EC 1.1.1.23. D-Galactarate is involved in reactions catalyzed by 4.2.1.158 that is present in *Oceanobacillus iheyensis* [52].

**Fig. 5:**
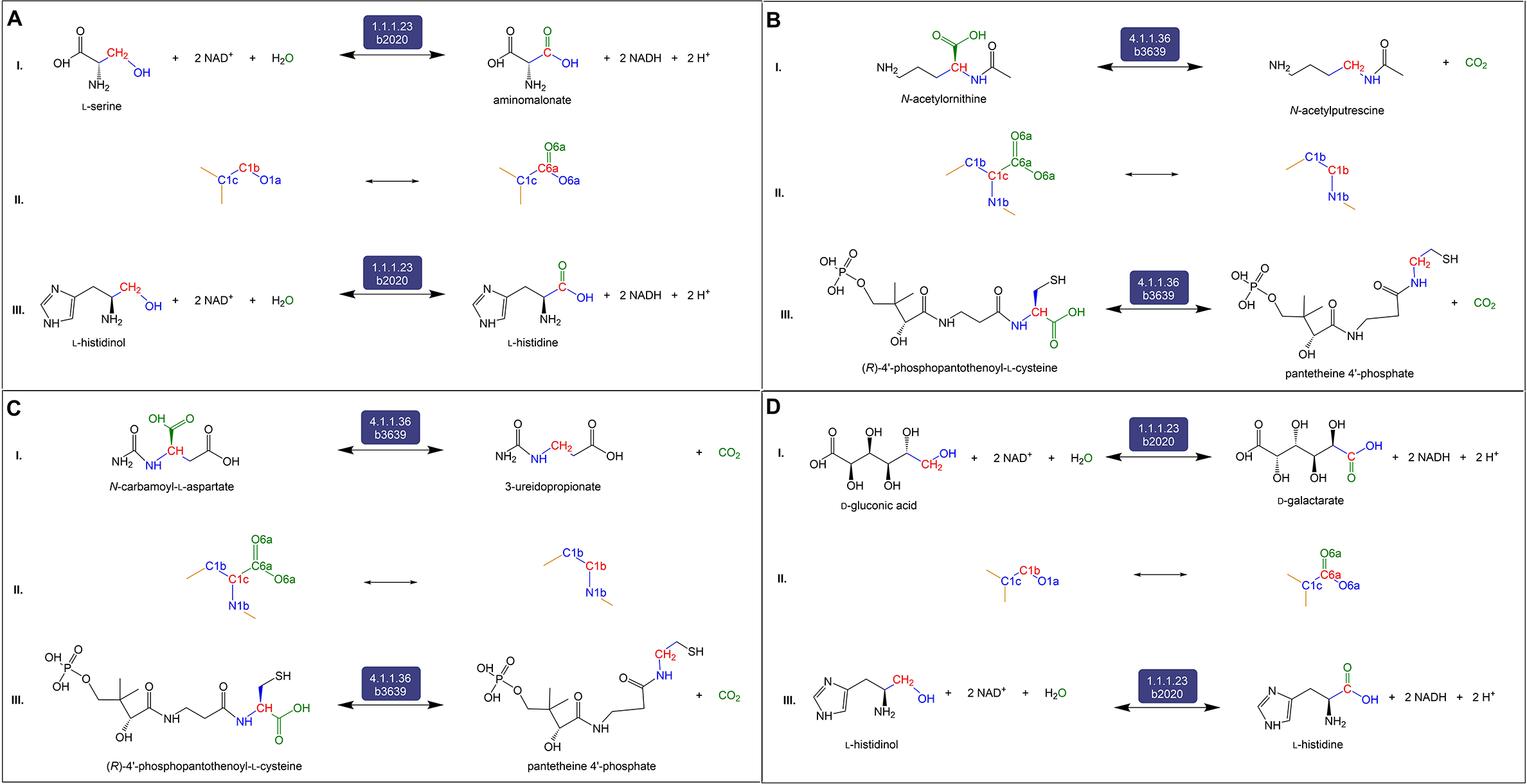
The set of four reactions belonging to Category 4 (C4). C4 reactions and derivatives are neither present in iML1515 nor associated with any other organism in KEGG or EcoCyc. Each of the four panels is divided into three sections I) the balanced reaction developed by our workflow indicating the reactants, products, and the promiscuous enzyme, II) the RDM pattern showing the Reaction Center (R) in red, and III) the native reaction catalyzed by the potentially promiscuous enzyme, as catalogued in KEGG.

## Discussion

Current practices for reconstructing genome-scale metabolic models, which are derived using sequencing and functional annotation, can be improved by utilizing metabolomics data. However, directly utilizing metabolomics measurements to augment existing models is challenging. Not every metabolite is measurable due to limited resolution and fidelity of mass spectrometry instruments. Further, assigning chemical identities to measured metabolites remains a challenge. Even if new metabolites are identified, their formation cannot be easily assigned to enzymes without significant experimental effort involving either genetic or biochemical screens. Additionally, metabolomics data alone cannot differentiate reactions catalyzed by different enzymes yet between the same substrates-product pairs without additional experimental efforts. Computational tools and workflows, as presented in this paper, can significantly guide such studies and aid in metabolic model construction and augmentation based on metabolomics data.

The workflow we developed here is designed to identify metabolites that can form due to promiscuous enzymatic activity within a specific model organism. Further, the workflow provides balanced reactions to document such enzymatic activities. We utilized PROXIMAL [39], which first identifies patterns of structural transformations associated with enzymes in the biological sample and then applies these transformations to known sample metabolites to predict putative metabolic products. Using PROXIMAL in this way allows attributing putative metabolic products to specific enzymatic activity and deriving balanced biochemical reactions that capture the promiscuous activity. Using PROXIMAL offers an additional advantage – the derived promiscuous transformations are specific to the sample under study and are not limited to hand-curated biotransformation operators as in prior works [33, 34]. PROXIMAL therefore allows exploration of a variety of biotransformations that are commensurate with the biochemical diversity of the biological sample. The EMMA workflow, which utilized PROXIMAL, was previously developed to engineer a candidate set from a metabolic model for metabolite identification [53]. EMMA did not aim to augment existing metabolic models or derive balanced reactions as utilized in this study.

Future experimental and computational efforts can further advance this work. Experimentally, the list of putative products generated by PROXIMAL but not documented in any metabolomics databases can be used as a resource to identify as yet unidentified metabolites. Experimental validation of reactions in the C1, C3 and C4 categories would provide further evidence of the suggested reactions, and would provide a means for expanding existing databases such as KEGG and EcoCyc. Computationally, PROXIMAL can be upgraded to consider enzymes that act on more than one Reaction Center (R) within a metabolite (e.g. transketolase). This would produce multiple operators per reaction and generate a more comprehensive list of putative reactions and products. When applying PROXIMAL, we did not consider whether products of promiscuous reactions can themselves act as new substrates for promiscuous reactions. This is due to the large number of putative products. We are currently developing machine learning techniques to improve the prediction accuracy of PROXIMAL.

## Conclusion

This study investigates creating Extended Metabolic Models (EMMs) through the augmentation of existing metabolic models with reactions due to promiscuous enzymatic activity. Our workflow, EMMA, first utilizes PROXIMAL to predict putative metabolic products, and then compares these products against metabolomics data. EMMA was applied to iML1515, the genome-scale model of *E. coli* MG1655. PROXIMAL generated 1,875 biochemical operators based on reactions in iML1515 and predicted 1,368 derivatives of 106 high-concentration metabolites. To validate these products, EMMA compared the set of putative derivatives with the set of metabolites documented in ECMDB as part of *E. coli* metabolism. For the overlapping set, we generated corresponding atom-balanced reactions by adding suitable cofactors and/or co-substrates to the substrate-derivative pair suggested by PROXIMAL. The balanced reactions were compared with data recorded in EcoCyc and KEGG. Our workflow generated a list of 23 new reactions that should be utilized to extend the iML1515 model, including parallel reactions between existing metabolites, novel routes to existing metabolites, and new paths to new metabolites. Importantly, this study is foundational in providing a systemic way of coupling computational predictions with metabolomics data to explore the complete metabolic repertoire of organisms. The described workflow can be applied to any organism utilizing its metabolic model to predict sample-specific promiscuous enzymatic byproducts. Applying this workflow to other biological samples and their metabolomics data promise to enhance our understanding of natural, synthetic, and xenobiotic metabolism.

## Methods

The EMMA workflow was customized to augment the *E. coli* iML1515 model based on the availability of the metabolic measurements in ECMDB, and the availability of cataloged reactions and metabolites for *E. coli* in other databases (EcoCyc and KEGG) (**Fig. 6**). The iML1515 model consists of 1,877 metabolites, 2,712 reactions and 1,516 genes. Our workflow consists of the following three steps.

**Fig. 6:**
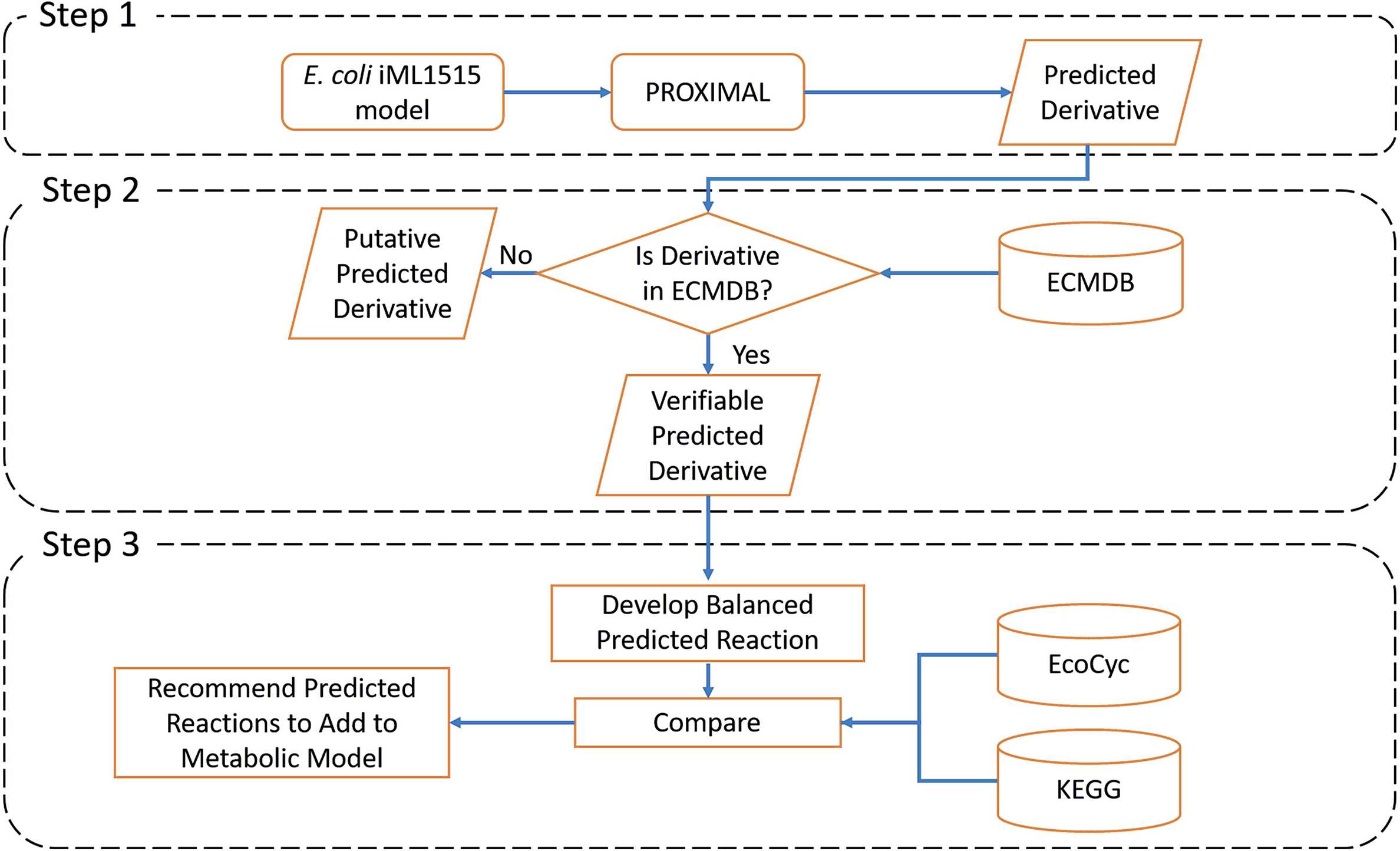
Main steps of EMMA workflow customized to extend the *E. coli* iML1515 model with predicted reactions. Step 1: Predict promiscuous transformations and derivatives using PROXIMAL. Step 2: Compare derivatives with measured metabolic dataset(s). Step 3: Curation and stoichiometric balancing of reactions.

### Step 1 – Predict promiscuous products using PROXIMAL

EMMA used PROXIMAL to predict putative products that can be added to the model. PROXIMAL utilizes RDM patterns [40] specific to the model’s reactions to create lookup tables that map reaction centers to structural transformation patterns. An RDM pattern specifies local regions of structural similarities/differences for reactant-product pairs based on a given biochemical reaction. An RDM pattern consists of three parts: i) A Reaction Center (R) atom exists in both the substrate and reactant molecule and is the center of the molecular transformation. ii) Difference Region (D) atoms are adjacent to the R atom and are distinct between substrate and product. iii) Matched Region (M) atoms are adjacent to the R atom but remain unmodified by the transformation. All atoms are labelled using KEGG atom types [54]. PROXIMAL constructs a lookup table of all possible biotransformations that can occur due to promiscuous activity of enzymes based on the RDM patterns of reactions catalyzed by enzymes associated with genes in the iML1515 gene list. The “key” in the lookup table consisted of the R and M atom(s) in the reactant, while the “value” is the R and D atom(s) in the product. RDM patterns were initially available through the (RPAIR) database, but they are now catalogued in KEGG’s RClass database. The biotransformation operators in the lookup table were then applied to model metabolites. The outcome of this step is a list of predicted products due to putative enzymatic activity.

### Step 2 – Compare predicted products with metabolomics dataset

Metabolites predicted by PROXIMAL were compared with measured metabolic data in ECMDB. ECMDB contains 3,760 metabolites detected in *E. coli* strain K-12 and related information such as reactions, enzymes, pathways, and other properties. This information was either collected from resources and databases such as EcoCyc, KEGG, EchoBase [55], UniProt [56, 57], YMDB [58], and CCDB [59], or from literature, or validated experimentally by the creators of ECMDB. Partial information about metabolites such as KEGG compound IDs, metabolites cell location, and chemical formulas is provided in ECMDB.

For each putative product, a mol file was generated and then converted to a SMILES string using Pybel [60], a python wrapper for the chemical toolbox Open Babel [61]. Based on the SMILES string, we initially retrieved the corresponding PubChem ID and InchiKey from PubChem using Pybel. To ensure consistency, we confirmed that retrieved PubChem IDs and InchiKeys of PROXIMAL predicted metabolites matched the corresponding entries in ECMDB. During this process, we noted some discrepancies. In some cases, the information retrieved from PubChem, such as InchiKeys did not match those in ECMDB. In cases of a mismatch, we sought additional information to confirm metabolite identities of ECMDB products. We utilized the values of the CAS ID, BioCyc ID, Chebi ID and KEGG ID fields to retrieve PubChem IDs using Pybel. The retrieved PubChem IDs are used to determine the ID through a majority vote. For example, if the PubChem ID associated with InchiKey, KEGG ID and CAS ID matched, but did not match the PubChem ID provided in ECMDB, then we considered the one retrieved by Pybel as the correct PubChem ID. Out of 3,760 metabolites in ECMDB, we identified 3,397 metabolites with consistent information with data retrieved from PubChem. Once PubChem IDs were identified for ECMDB metabolites, we compared our predicted metabolites against ECMDB metabolites using PubChem IDs.

### Step 3 – Curation of stoichiometric reactions

If a metabolite predicted by PROXIMAL was in ECMDB, then steps 1 and 2 resulted in the identification of a *verifiable* predicted promiscuous transformation of an *E. coli* metabolite. Each predicted transformation was manually examined and compared against the RDM pattern causing the transformation. Transformations were discarded if the they seemed infeasible, if the substrate was a cofactor, or if the RPAIR entry associated with the PROXIMAL operator required the presence of more than one Reaction Center (R). For each valid verifiable predicted transformation by PROXIMAL, we developed a new reaction by examining the reaction(s) template associated with the enzymatic transformation and adding suitable cofactors to the reactant and product of the biotransformation identified. The set of developed balanced reactions, where the added cofactors to a reaction caused the number of atoms of reactants and products to match on both sides of the reaction, are then compared to reactions recorded in EcoCyc, KEGG, or the literature.

The outcomes were divided into four categories. C1 reactions consisted of metabolites predicted by PROXIMAL that are already in iML1515 but catalyzed by different enzymes than the ones already listed in the model. These reactions reflect promiscuous activity that enabled the same biotransformation catalyzed by a different gene in the model. C2 reactions already existed in EcoCyc and/or KEGG but not in iML1515. This reflected a curation problem where some reactions were not included in the iML1515 model. C3 reactions were not in EcoCyc but documented in KEGG for other organisms. C4 reactions did not exist in either EcoCyc nor in KEGG. These reactions were thus novel reactions that have not been reported in the literature.

## Declarations

### Ethics approval and consent to participate

Not applicable

### Consent for publication

Not applicable

### Availability of data and material

The *E. coli* iML1615 model is available on the BiGG database and can be found at http://bigg.ucsd.edu/models/iML1515. Data from ECMDB can be directly downloaded from the ECMDB website. A full list of derivatives that were predicted by PROXIMAL and had a chemical ID in PubChem can be found in **Supplementary File 1**.

### Author’s contribution

SH conceived the EMMA concept. SA developed the EMMA workflow. EC curated the results. VP verified the analysis. NN and SH supervised the work done through the development of the workflow and data curation. Manuscript was written by SA and EC, reviewed by VP, and revised by NN and SH.

## Acknowledgments

Not applicable

