## Supplementary File 2 for "Towards creating an extended metabolic model (EMM) for *E. coli* using enzyme promiscuity prediction and metabolomics data"

### Towards creating an extended metabolic model (EMM) for *E. coli* using enzyme promiscuity prediction and metabolomics data

Sara A. Amin, Elizabeth Chavez, Vladimir Porokhin, Nikhil U. Nair, and Soha Hassoun

#### Evaluating contributions of new reactions to biomass production

The EMMA workflow suggested 23 reactions that can be utilized to extend the *E. coli* iML1515 model. We first performed Flux Balance Analysis (FBA) to assess the impact on biomass production when augmenting iML1515 with each of the EMMA reactions. The maximum flux value of the biomass reaction before adding EMMA reactions was 0.877 mmol/g CDW/h. We then in turn added each EMMA reaction and assessed the change in biomass flux. There were no changes in the maximum flux value.

Next, we performed Flux Variability Analysis (FVA) of the added EMMA reactions. The biomass flux was fixed at 0.877 mmol/g CDW/h. The flux through each EMMA reaction was then in turn maximized and minimized under this biomass constraint. Table S2 summarizes our findings. Each row is associated with a substrate (first column), a predicted product (second column) derived using PROXIMAL, and the developed EMMA reaction (third column). The minimum (fourth column) and maximum fluxes (fifth column) of each EMMA reaction are reported. FVA shows that the minimum flux value was zero for all reactions. A few reactions had zero maximum flux value, indicating that the EMMA reaction cannot carry any flux. Other reactions had a small (in magnitude) maximum flux value. The directionality of the maximum flux varied, indicating that the predicted transformation is reversible.

**Table S2:** FVA analysis results when adding each of the EMMA reactions (last column) to *E. coli* iML1515.

| Substrate | Predicted Product | Reaction | Min Reaction Flux (mmol/g CDW/h) | Max Reaction Flux (mmol/g CDW/h) |
|---|---|---|---|---|
| Pyruvate | 4-Hydroxy-2-oxoglutarate | Pyruvate + Glyoxylate ⟷4-Hydroxy-2-oxoglutarate | 0 | 9.83E-13 |
| Pyruvate | 4-Carboxy-4-hydroxy-2-oxoadipate | Pyruvate + Oxaloacetate ⟷ 4-Carboxy-4-hydroxy-2-oxoadipate | 0 | -9.71E-13 |
| L-Glutamate | γ-Glutamyl-β-cyanoalanine | L-Glutamate + 3-Cyano-L-alanine ⟷ γgamma-Glutamyl-βbeta-cyanoalanine + $H_2O$ | 0 | 0 |
| L-Glutamate | THF-L-glutamate | ATP + THF + L-Glutamate ⟷ ADP + Phosphate + THF-L-glutamate | 0 | -1.25E-13 |
| 2-Oxoglutarate | 2-Hydroxyglutarate | 2-Oxoglutarate + NADPH + H+ ⟷ 2-Hydroxyglutarate + NADP+ | 0 | -4.32E-12 |
| 2-Oxoglutarate | 2-Oxoglutaramate | 2-Oxoglutarate + Ammonia ⟷ 2-Oxoglutaramate + $H_2O$ | 0 | 0.00E+00 |
| L-Alanine | Dehydroalanine | L-Alanine + NADP+ ⟷ Dehydroalanine + NADPH + H+ | 0 | 8.70E-12 |
| L-Serine | Aminomalonate | L-Serine + 2 NAD+ + $H_2O$ ⟷ Aminomalonate + 2 NADH + 2 H+ | 0 | 1.15E-11 |
| Propanoyl-CoA | 2-Methylacetoacetyl-CoA | Propanoyl-CoA + Acetyl-CoA ⟷CoA + 2-Methylacetoacetyl-CoA | 0 | 3.90E-13 |
| L-Histidine | Imidazol-5-yl-pyruvate | L-Histidine + 2-Oxoglutarate ⟷ Imidazol-5-yl-pyruvate + L-Glutamate | 0 | 4.97E-13 |
| 4-Hydroxybenzoate | Geranyl-hydroxybenzoate | 4-Hydroxybenzoate + Geranyl diphosphate ⟷ Geranyl-hydroxybenzoate + Diphosphate | 0 | 6.45E-13 |
| Phenylpyruvate | Phenyllactate | Phenylpyruvate + NADPH + H+ ⟷ Phenyllactate + NADP+ | 0 | 2.62E-13 |
| D-Ribulose 5-phosphate | D-Ribulose 1,5-bisphosphate | ATP + D-Ribulose 5-phosphate ⟷ ADP + D-Ribulose 1,5-bisphosphate | 0 | -2.56E-13 |
| D-Gluconic acid | 2-Keto-D-gluconic acid | D-Gluconic acid + NADP+ ⟷ 2-Keto-D-gluconic acid + NADPH + H+ | 0 | -2.01E-14 |
| D-Gluconic acid | D-Galactarate | D-Gluconic acid + + 2 NAD+ + $H_2O$ ⟷ D-Galactarate + 2 NADH + 2 H+ | 0 | -2.78E-12 |
| D-Glycerate | Tartrate | D-Glycerate + $CO_2$ ⟷ (*R,R*)-Tartaric acid | 0 | 3.58E-12 |
| Bicarbonate | Carboxyphosphate | Bicarbonate + ATP ⟷ Carboxyphosphate + ADP | 0 | -4.16E-11 |
| Bicarbonate | Carboxyphosphate | Bicarbonate + ATP ⟷ Carboxyphosphate + ADP | 0 | -4.16E-11 |
| Acetoacetyl-CoA | Acetoacetate | Acetoacetyl-CoA + Succinate ⟷ Acetoacetate + Succinyl-CoA | 0 | 2.53E-12 |
| Cytosine | CMP | CMP + $H_2O$ ⟷ Cytosine + D-Ribose 5-phosphate | 0 | 1.19E-12 |

| | | | | |
|---|---|---|---|---|
| N-Acetylornithine | N-Acetylputrescine | N-Acetylornithine ↔ *N*-Acetylputrescine + $CO_2$ | 0 | -4.73E-12 |
| N-Carbamoyl-L-aspartate | 3-Ureidopropionate | *N*-Carbamoyl-L-aspartate ↔ 3-Ureidopropionate + $CO_2$ | 0 | 5.51E-12 |
| 1-(5'-Phosphoribosyl)-5-amino-4-imidazolecarboxamide | 5-Amino-4-imidazolecarboxyamide | 1-(5'-Phosphoribosyl)-5-amino-4-imidazolecarboxamide + Diphosphate ↔ 5-Amino-4-imidazolecarboxyamide + 5-Phospho-alpha-D-ribose 1-diphosphate | 0 | 4.92E-14 |
